# An RNase bacterial toxin family cleaves the anticodon loop across diverse tRNA isotypes

**DOI:** 10.64898/2026.11.23.768618

**Authors:** Jacob E. Lin, Audrey R. Long, Kristi Pham, Ethel Bayer-Santos

## Abstract

Characterized bacterial tRNase toxins typically recognize individual tRNAs or restricted tRNA subsets. Here, we identify TloR (tRNA loop-targeting RNase), a tRNase with an unusually broad substrate range but a strong positional preference for exposed tRNA loops. Comparative genomic analyses identified TloR homologs across diverse bacterial lineages, where they are frequently embedded in conflict-associated loci. Structural modeling supports assignment of TloR to the recently described BECR-like superfamily, and mutational analyses identify conserved histidines required for toxicity and RNA cleavage. TloR cleaved tRNA and inhibited protein synthesis in a reconstituted translation system without detectable degradation of ribosomal RNA or plasmid DNA. Small-RNA sequencing showed that toxin-induced fragments arise from tRNAs decoding 19 of the 20 amino acids, with fragment endpoints strongly enriched near the anticodon loop. Purified TloR directly cleaved an unmodified tRNA substrate, predominantly within the anticodon loop. TloR therefore defines a family of BECR-like tRNases that combines broad tRNA recognition with positional preference for exposed tRNA loops. Together, these findings establish broad but positionally constrained tRNA cleavage as a distinct targeting strategy for antibacterial RNases.

## INTRODUCTION

Microorganisms deploy polymorphic toxins to disable competitors and reshape access to nutrients and shared niches [1, 2]. These modular proteins couple conserved trafficking regions to rapidly diversifying C-terminal toxin domains, allowing distinct toxic activities to be exchanged among delivery systems. Immunity proteins encoded adjacent to each toxin gene protect toxin-producing cells and their sister cells from self-intoxication [3]. Ribonuclease toxins are particularly effective because cleavage of even a small number of critical RNA molecules can disrupt essential information-processing pathways. Among them, tRNA-targeting ribonucleases rapidly inhibit translation by depleting functional tRNAs [4–6].

Most characterized bacterial tRNase toxins exhibit restricted substrate recognition. Colicin D cleaves four arginine isoacceptors, whereas colicin E5 recognizes four tRNAs that share a QUN anticodon sequence [7, 8]. VapC, MazF, and contact-dependent inhibition (CDI) toxins likewise target individual tRNAs or defined subsets through combinations of sequence, structure, modification state, and translation-factor dependence [9–12]. Thus, substrate discrimination among tRNAs is a recurring feature of bacterial tRNase activity. Broad tRNA recognition coupled to positional cleavage specificity remains largely unexplored among polymorphic toxins.

The BECR fold, named for Barnase, EndoU, Colicin E5/D, and RelE, has repeatedly diversified into toxic ribonuclease domains [1, 13, 14]. Its compact α/β architecture supports extensive sequence and topological variation while preserving catalytic features compatible with metal-independent RNA cleavage. A recent structure-guided analysis expanded the known BECR landscape and identified a divergent BECR-like superfamily with similar catalytic organization but distinct topology and weak sequence similarity to canonical BECR proteins [14]. SRLase1, recently characterized by our group, established an RNA-targeting activity within this divergent superfamily. SRLase1 cleaves the sarcin-ricin loop of 23S rRNA through an unusual EF-Tu-dependent mechanism [15]. The substrate range and biological activities of other BECR-like proteins remain largely unexplored.

Here, we characterize TloR (tRNA loop-targeting RNase) and its cognate immunity protein, TloRI. By integrating comparative genomics, structural modeling, mutagenesis, biochemical assays, and transcriptome-wide mapping of cleavage-associated products, we show that TloR defines a widespread BECR-like toxin family that acts on tRNAs. Unlike most characterized tRNase toxins, TloR acts across a broad range of tRNA species while preferentially cleaving exposed tRNA loops, particularly the anticodon loop. These findings expand the known functional repertoire of the BECR-like superfamily and identify broad but positionally constrained tRNA cleavage as a distinct targeting strategy for antibacterial ribonucleases.

## RESULTS

### TloR and TloRI form a broadly distributed toxin-immunity pair

TloR (previously designated STox11) was initially identified as a predicted toxin domain encoded within a secretion-associated locus in *Salmonella enterica* [16]. To determine whether TloR represents a broader toxin family, we performed iterative searches for homologous proteins followed by manual sequence curation and structural validation using AlphaFold3 (AF3) [17]. These analyses identified 191 TloR homologs distributed across diverse bacterial lineages, including members of the Pseudomonadota, Bacillota, Bacteroidota, Cyanobacteriota, and Actinomycetota (Fig. 1A, Table S1). In addition, a single homolog was detected in a *Methanobrevibacter* genome (WP_320629730.1), indicating that although the family is predominantly bacterial, it also occurs in archaea (Fig. 1A). These observations suggest that TloR defines a widespread protein family with a broad phylogenetic distribution.

**Fig 1.**
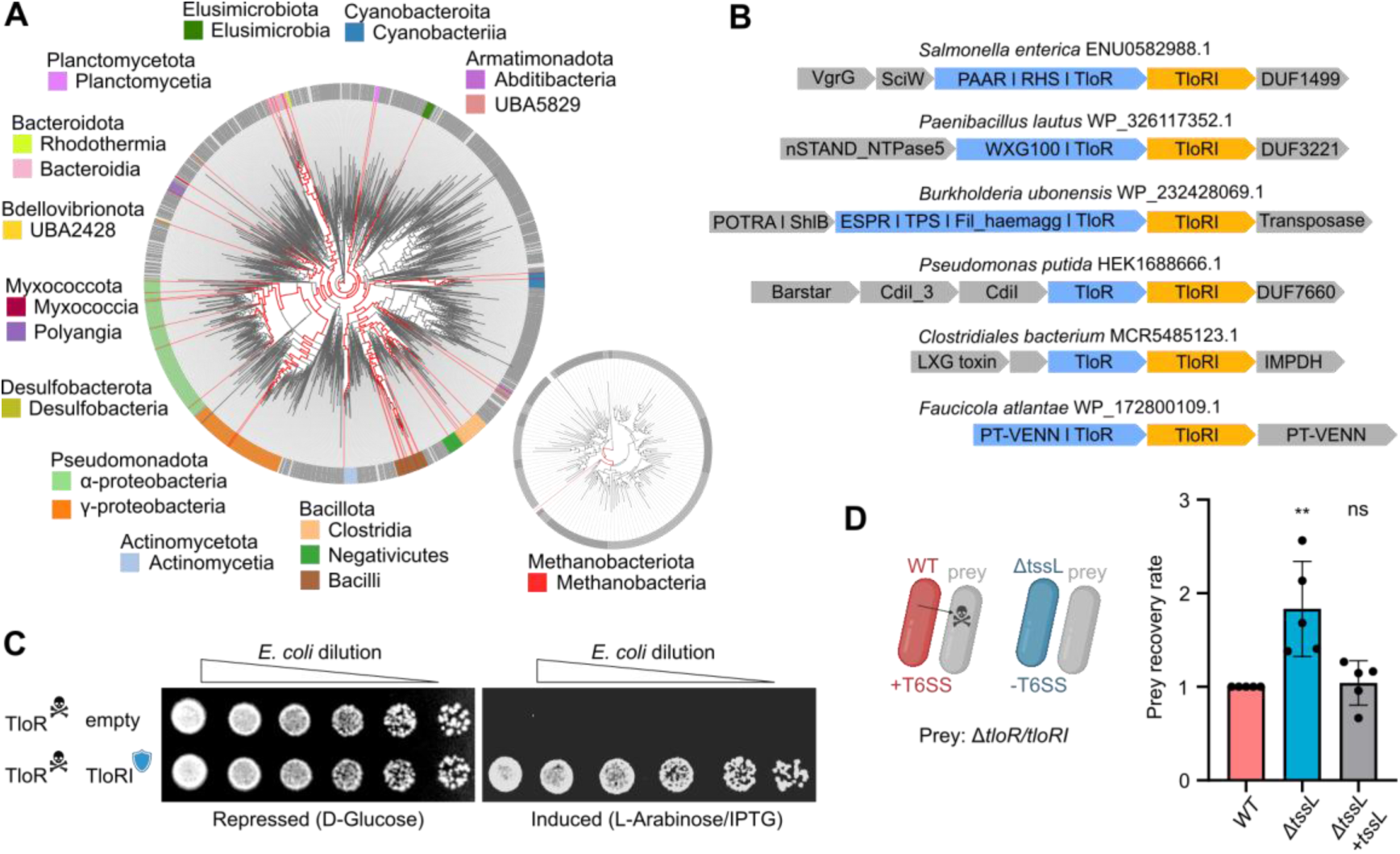
TloR and TloRI form a widespread toxin-immunity pair associated with several conflict systems. **(a)** Phylogenetic tree showing the distribution of TloR throughout various bacteria and archaea. Classes containing TloR labelled on the outer ring are listed under their respective phyla. **(b)** Genomic organization of representatives showing TloR and TloRI encoded in the context of conflict systems. **(c)** *E. coli* toxicity assay. Serial dilutions of *E. coli* carrying TloR and TloRI constructs plated in repressive or inductive conditions. Images are representative of three independent experiments. **(d)** Interbacterial competition assay. Wild-type *S.* Oranienburg outcompetes prey while the deletion of T6SS component *tssL* eliminates this competitive advantage. Data represents the mean ± SD of five independent experiments. Statistical significance was assessed by one-way ANOVA followed by Dunnett’s multiple comparison test. \*\**p*<0.01.

To gain insight into the biological role of TloR, we examined the genomic neighborhoods of all 191 homologs (Dataset S1). TloR homologs were frequently embedded within conflict-associated genomic loci characteristic of polymorphic toxin systems. In Gram-negative bacteria, TloR was commonly associated with CDI systems [18] and T6SS [19], often occurring as the C-terminal toxin domain of RHS proteins [20] or fused to PAAR-containing delivery modules [21]. In Gram-positive bacteria, TloR was associated with loci encoding components of the Type VII secretion system (T7SS) [22]. Together, these genomic architectures place TloR within multiple secretion systems used during interbacterial antagonism and strongly support its role as an antibacterial effector (Fig. 1B).

Most TloR homologs are followed by a small gene predicted to encode a cognate immunity protein. A sequence similarity network of these candidate immunity proteins identified two major clusters (Fig. S1A). The larger cluster contained domains related to Imm55, a family defined previously by computational analysis of polymorphic toxin systems [1]. This cluster retained detectable sequence similarity to Imm55 and shared its predicted structural features but resolved as a separate group in the network, and we therefore designate it Imm55.2 (Fig. S1A). The second cluster showed no detectable similarity to Imm55 or Imm55.2 and defines a new immunity protein family, which we name Imm70 following the nomenclature proposed by Zhang et al. (2012). Imm70 and Imm55.2 share little sequence or structural similarity with each other, even though the TloR domains they neutralize are structurally conserved (Fig. S1B,C). TloR toxins therefore pair with at least two unrelated immunity protein families.

To experimentally determine whether TloR and its cognate immunity protein constitute a functional toxin-immunity pair, we selected a representative locus from the T6SS of *Salmonella enterica* serovar Oranienburg (ENU0582988.1) and co-expressed them in *Escherichia coli*. Expression of TloR strongly inhibited bacterial growth, whereas co-expression of its cognate immunity protein TloRI, which carries an Imm55.2 domain, fully restored growth (Fig. 1C). These results establish TloR and TloRI as a bona fide toxin-immunity pair. We next tested whether TloR contributes to T6SS-dependent competition in *S.* Oranienburg. Wild-type attackers reduced recovery of the Δ*tloR/tloRI* prey strain, whereas deletion of *tssL*, which encodes an essential T6SS component, eliminated this competitive advantage (Fig. 1D). Complementation of *tssL* restored the competitive advantage of the attacker. TloR is therefore deployed through the T6SS during interbacterial competition. Together with the broad phylogenetic distribution of TloR and its recurrent association with conflict-related secretion systems and adjacent immunity genes, these findings support TloR as a widespread family of antibacterial polymorphic toxins.

### TloR defines a BECR-like toxin family with a conserved catalytic pocket and immunity-binding interface

To gain insight into TloR function, we analyzed AF3 models together with sequence conservation across homologs. TloR adopts the characteristic BECR-like architecture, consisting of an α-helix followed by a series of meandering β-strands that form the conserved structural core of the recently described BECR-like superfamily (Fig. 2A) [14]. Despite this conserved core, TloR contains a distinct arrangement of peripheral structural elements. Most BECR-like proteins contain an additional β-strand preceding the central α-helix, whereas TloR lacks this element and instead contains a β6 strand in the corresponding region (Fig. 2A). These features distinguish TloR as a structurally distinct family within the broader BECR-like superfamily.

**Fig 2.**
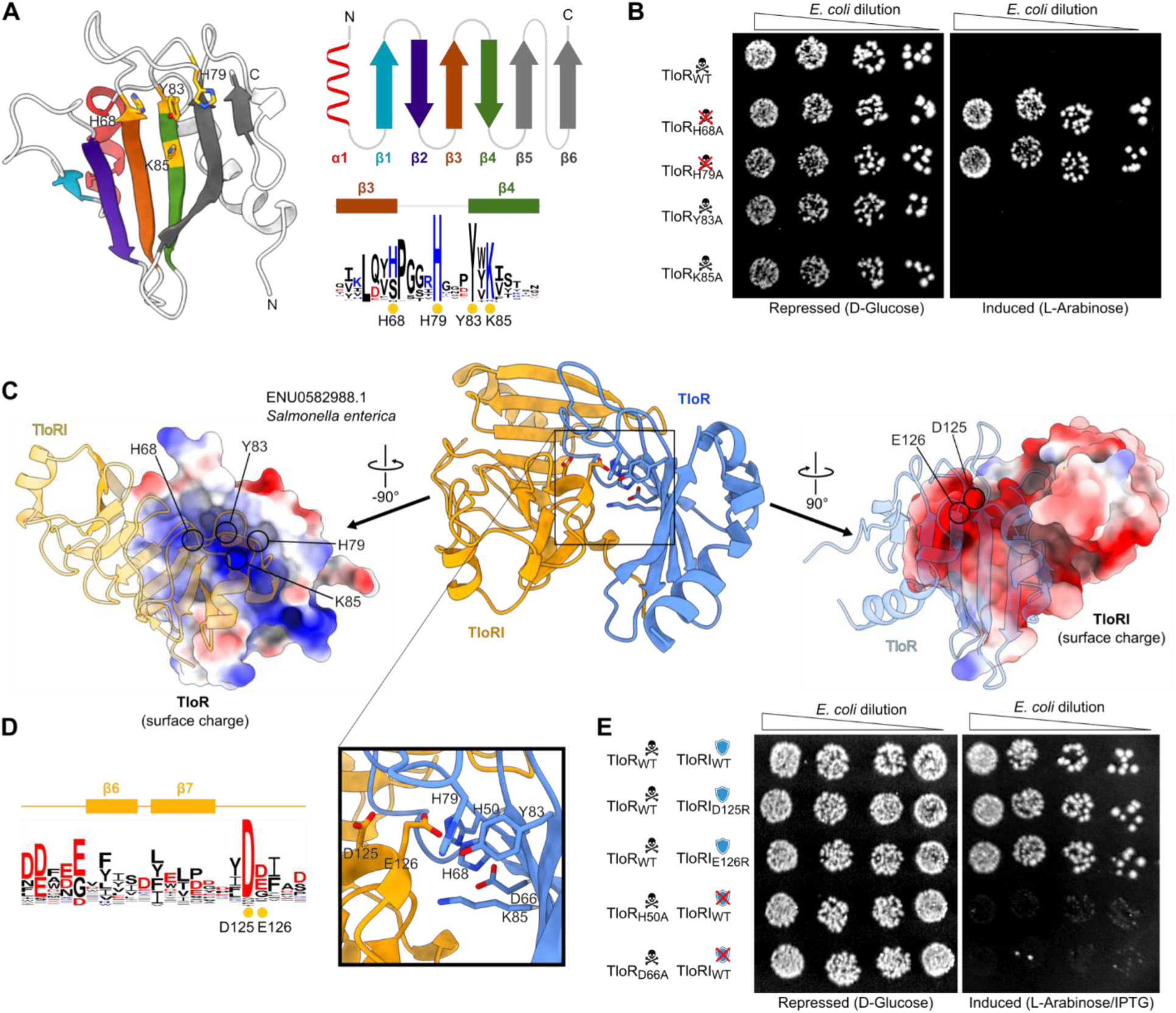
TloR adopts a BECR-like fold with a predicted active-site occluded by TloRI. **(a)** AF3 model of TloR. Yellow residues indicate amino acids analyzed by mutagenesis. TloR sequence logo is colored by amino acid charge, with secondary structure and mutated residues indicated. **(b)** TloR toxicity assay. Serial dilutions of *E. coli* carrying wild-type or mutant TloR were plated under repressive or inductive conditions. Images are representative of three independent experiments. **(c)** AF3 model of the TloR-TloRI complex. Conserved TloR residues are shown at the interface with TloRI (left), and conserved TloRI residues are shown at the interface with TloR (right). **(d)** TloRI sequence logo colored by amino acid charge (left) and predicted toxin-immunity interface with putative interacting residues shown as sticks (right). Secondary structure and mutated residues are indicated. **(e)** TloR-TloRI neutralization assay. Serial dilutions of *E. coli* carrying the indicated wild-type or mutant TloR and TloRI variants were plated under repressive or inductive conditions. Images are representative of three independent experiments.

To identify residues that may contribute to TloR activity, we mapped sequence conservation onto the predicted structure and examined conserved residues within the largest positively charged surface pocket (Fig. 2A). This analysis identified a highly conserved HxxxYxK motif comprising His79, Tyr83, and Lys85, together with a second conserved histidine, His68. These residues cluster within the positively charged pocket and are among the most conserved residues across TloR homologs. The composition of this putative active site, which is enriched in aromatic and basic residues, resembles catalytic sites of previously characterized BECR ribonucleases [7, 13, 23]. Its pronounced positive electrostatic potential is also consistent with binding to the negatively charged phosphate backbone of nucleic acid substrates.

We first asked which of these conserved residues are required for toxin activity. Alanine substitution of either His68 or His79 abolished TloR-mediated toxicity in *E. coli*, whereas Y83A and K85A remained toxic (Fig. 2B). Western blot analysis confirmed detectable accumulation of the H79A variant (Fig. S2). These results identify His68 and His79 as residues required for TloR activity and, together with their conservation and location within the predicted active-site pocket, support their involvement in catalysis.

We next asked whether residues surrounding this pocket contribute to recognition by the cognate immunity protein TloRI. The AF3 model of the TloR-TloRI complex predicts that a negatively charged surface of TloRI binds across the positively charged surface of TloR, positioning TloRI over the putative active-site (Fig. 2C,D). Two TloR residues at this predicted interface, His50 and Asp66, are predicted to contact the conserved TloRI residues Asp125 and Glu126 (Fig. 2D). To distinguish effects on toxin activity from effects on immunity recognition, we first tested whether H50A and D66A remained toxic in the absence of TloRI. Both variants retained toxicity comparable to wild-type TloR (Fig. S2), and western blot analysis confirmed their accumulation. We then tested whether wild-type TloRI could neutralize these active toxin variants. TloRI efficiently neutralized wild-type TloR but failed to protect cells from either H50A or D66A (Fig. 2E). Thus, His50 and Asp66 are dispensable for toxin activity but required for efficient neutralization by TloRI, supporting a specific role in toxin-immunity recognition. We also tested the reciprocal side of the predicted interface by substituting TloRI residues Asp125 and Glu126 with arginine. Neither D125R nor E126R alone abolished TloRI-mediated neutralization of wild-type TloR (Fig. 2E). Thus, disruption of a single predicted contact on the TloRI surface is insufficient to prevent neutralization, consistent with an interface supported by multiple interactions [24].

Together, these results support a model in which a single positively charged surface of TloR mediates both substrate recognition and immunity protein binding. His68 and His79 within this pocket are required for toxin activity, whereas His50 and Asp66 contribute to recognition by TloRI without being required for toxicity. The negatively charged surface of TloRI likely mimics the RNA substrate, occupying the substrate-binding surface and occluding the putative active site.

### TloR is an RNase that inhibits protein synthesis

Although members of the canonical BECR superfamily predominantly function as ribonucleases, the biochemical activities of BECR-like proteins remain largely unexplored. To define the molecular target of TloR, we examined whether toxin expression affected either DNA or RNA integrity. TloR expression did not activate the P*_recA_*::GFP SOS-response reporter, whereas the DNase toxin TseV2 [25] strongly induced the reporter (Fig. 3A). Consistent with this result, no degradation of plasmid DNA was detected following TloR expression in *E. coli* or after incubation of purified TloR with plasmid DNA *in vitro* (Fig. 3B,C; Fig. S3). These results argue against DNA as the primary cellular target of TloR.

**Fig 3.**
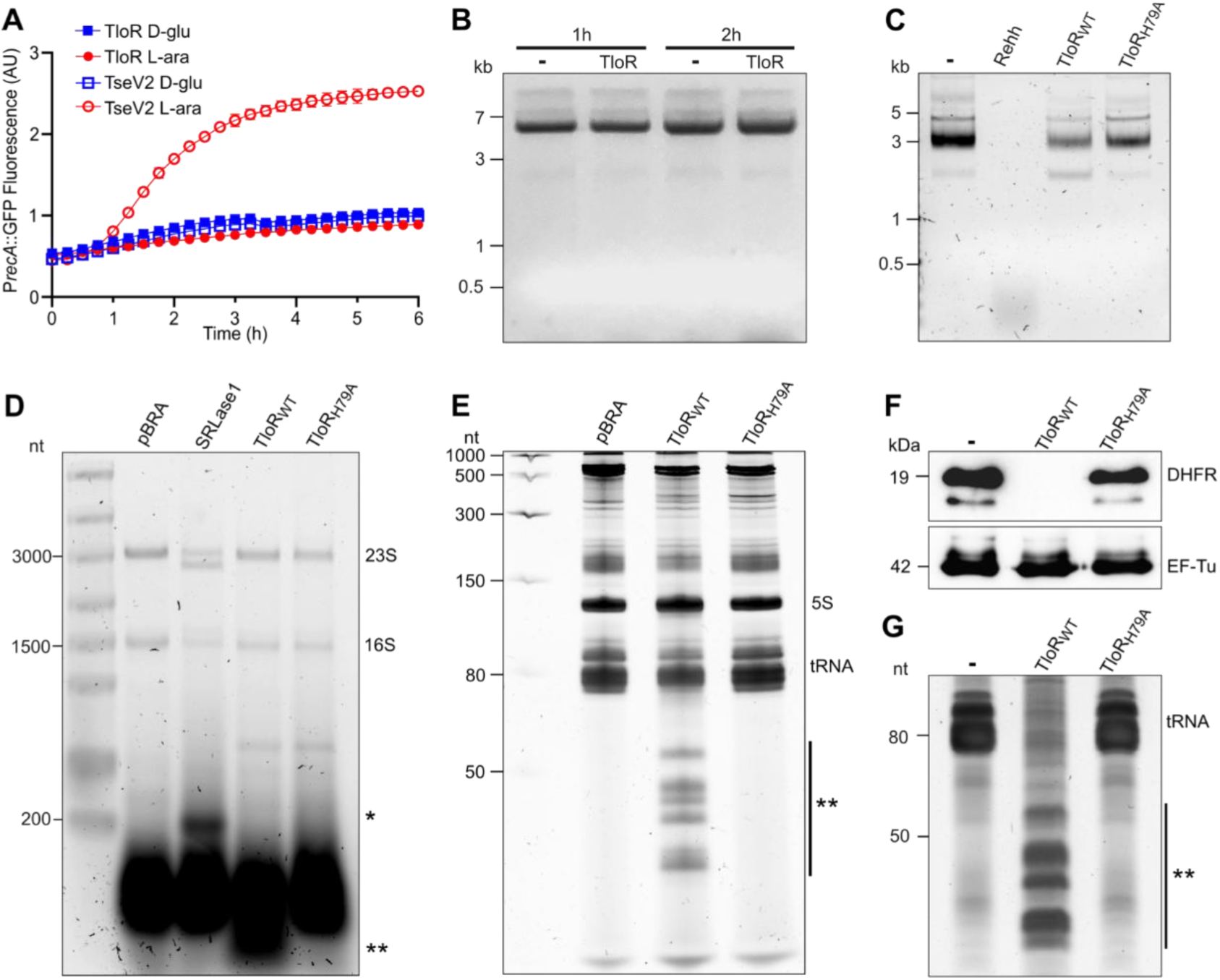
TloR cleaves RNAs and inhibits translation. **(a)** SOS-response reporter assay in *E. coli* expressing TloR. The DNase TseV2 was used as a positive control. Data represent the mean ± SD of three independent experiments. **(b, c)** Plasmid DNA degradation assays following TloR expression in *E. coli* **(b)** or incubation with purified TloR *in vitro* **(c)**. The DNase Rehh was used as a positive control in (c). **(d, e)** Analysis of total RNA isolated from *E. coli* expressing TloR_WT_, TloR_H79A_, or empty vector. RNA was resolved by agarose gel electrophoresis to visualize rRNAs **(d)** or denaturing urea-PAGE to resolve small RNAs **(e)**. SRLase1 was used as a positive control for rRNA cleavage (d). Asterisks indicate rRNA cleavage (*) or small RNA (**) products. **(f, g)** Effect of purified TloR on cell-free protein synthesis. TloR_WT_ or TloR_H79A_ was added to PURExpress reactions, and reporter protein synthesis was analyzed via western blot **(f)**. EF-Tu served as a loading control. RNA from the same reactions was analyzed by denaturing urea-PAGE **(g)**.

We next examined total RNA isolated from *E. coli* expressing TloR_WT_, the inactive TloR_H79A_, or an empty-vector control. Agarose gel electrophoresis revealed no detectable degradation of 16S or 23S ribosomal RNA following TloR_WT_ expression (Fig. 3D); distinct from the pattern observed for SRLase1 [15]. In contrast, additional low-molecular-weight RNA species accumulated in cells expressing TloR_WT_ (** in Fig. 3D). Resolution of the same RNA samples by denaturing urea-PAGE revealed multiple fragments of approximately 30 to 60 nt that were absent from cells expressing TloR_H79A_ or the empty-vector control (Fig. 3E). The 5S rRNA band remained largely unchanged. Thus, TloR activity produces discrete small-RNA fragments rather than widespread degradation of cellular RNA.

We next asked whether TloR-dependent RNA cleavage was sufficient to inhibit protein synthesis. Addition of purified TloR_WT_ to a cell-free translation system strongly reduced synthesis of the reporter protein DHFR, whereas TloR_H79A_ had no detectable effect (Fig. 3F). RNA isolated from the same reactions showed accumulation of cleavage products resembling the same band pattern generated during TloR expression in *E. coli* (Fig. 3G). Together, these results establish TloR as an RNase whose activity generates discrete small-RNA fragments and inhibits protein synthesis.

### TloR broadly targets tRNAs and preferentially cleaves the anticodon loop

To identify the small RNA fragments generated during TloR intoxication, we sequenced 14-to 70-nt RNAs isolated from *E. coli* expressing TloR or an empty-vector control. TloR-enriched fragments mapped predominantly to tRNA loci (Fig. 4A; Table S2), identifying tRNAs as the major source of the cleavage products observed by urea-PAGE (Fig. 3E). These fragments arose from numerous tRNA isotypes with diverse anticodon sequences and collectively represented tRNAs decoding 19 of the 20 amino acids. Thus, unlike most characterized toxin tRNases, TloR acts broadly across the cellular tRNA pool.

**Fig 4.**
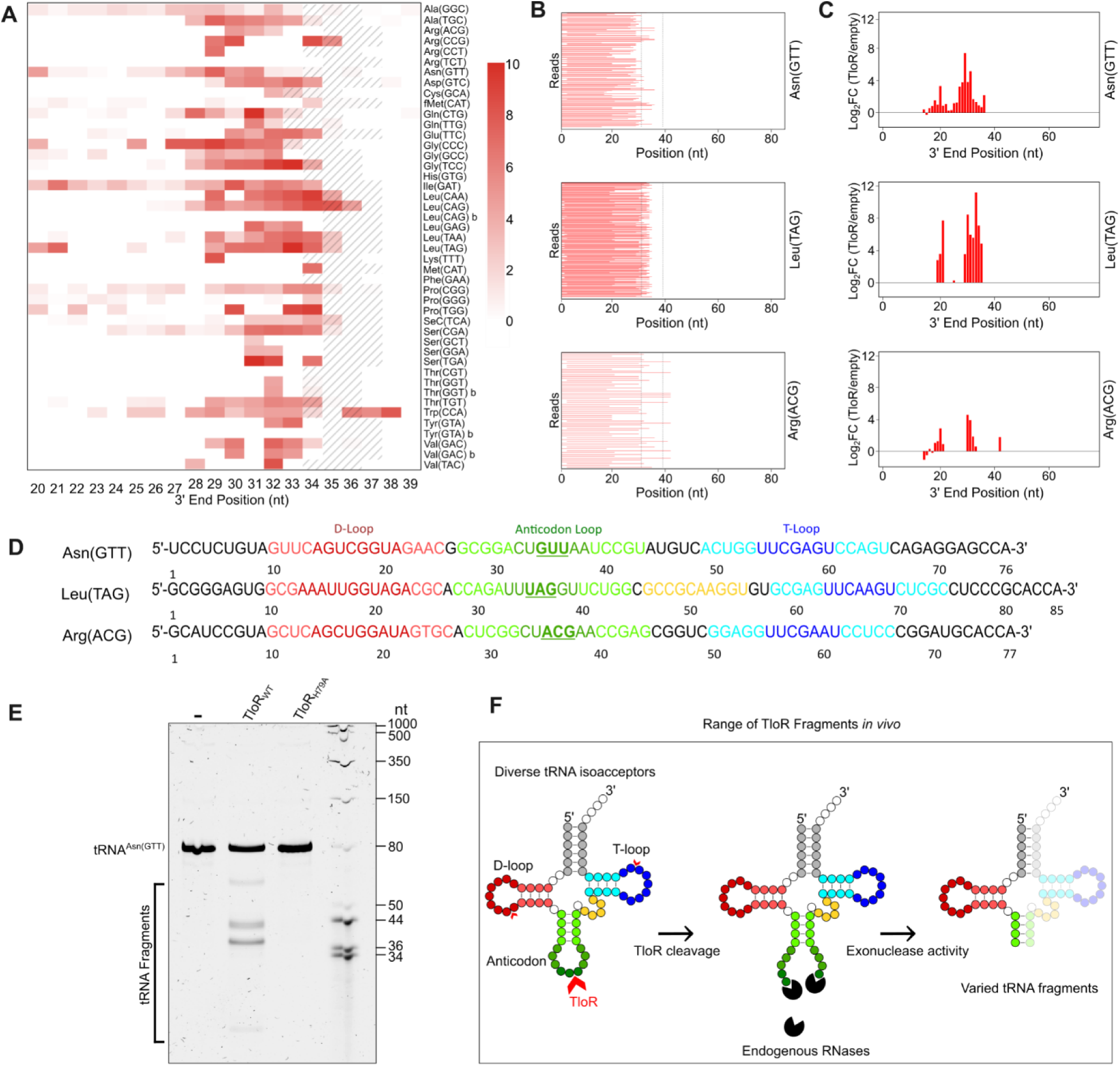
TloR cleaves diverse tRNA with a preference for the anticodon loop. **(a)** Distribution of TloR-enriched 5′ tRNA fragments. Heat map shows the log2 fold change (log2FC) in fragments from TloR-expressing *E. coli* relative to the empty-vector, plotted by tRNA species and 3′ fragment endpoint. Anticodon positions are indicated by hash marks. **(b)** Sequencing reads mapping to representative substrates. Read abundance is represented as the log2FC in TloR-expressing cells relative to the control. Dotted lines delimit the anticodon loop. **(c)** Endpoint distribution of TloR-enriched 5′ fragments from the representative tRNAs shown in (b). **(d)** Representative tRNA sequences. **(e)** Direct cleavage of *in vitro*-transcribed tRNA^Asn(GTT)^ by purified TloR. **(f)** Model relating direct TloR cleavage to tRNA fragments recovered from cells. TloR preferentially cleaves within the anticodon loop, with lower-frequency cleavage in other exposed loops. Subsequent processing by endogenous RNases is proposed to generate the heterogeneous 5′ fragment endpoints recovered by RNA sequencing.

We next examined where TloR-associated fragments accumulated within tRNAs sequences. For enriched 5′-derived fragments, we plotted their 3′ endpoints using mature tRNA coordinates (Fig. 4A). Across most tRNAs, these endpoints clustered predominantly at Sprinzl positions 28–34, spanning the distal anticodon stem and 5′ portion of the anticodon loop. Lower-abundance endpoints were also detected in other regions, including the D-loop. Representative sequence-divergent substrates, including tRNA^Asn(GTT)^, tRNA^Leu(TAG)^, and tRNA^Arg(ACG)^, showed similar endpoint distributions despite substantial differences in nucleotide sequence (Fig. 4B,C). Mapping the enriched fragments onto tRNA secondary structures further showed that fragment accumulation was concentrated near exposed loop regions rather than distributed uniformly across the molecule (Fig. 4D; Fig. S4). This data indicates that TloR-generated fragments exhibit a shared positional bias across sequence-divergent tRNAs.

Because cellular RNA fragments can undergo processing after toxin-mediated cleavage, we next determined the cleavage pattern produced directly by TloR. An unmodified *in vitro*-transcribed tRNA^Asn(GTT)^ was incubated with purified TloR_WT_ or TloR_H79A_ (Fig. S3) and analyzed by denaturing urea-PAGE (Fig. 4E). TloR_WT_ generated predominant cleavage products consistent with cleavage within the anticodon loop, whereas TloR_H79A_ showed no detectable activity. Lower-abundance products were also consistent with cleavage within other exposed loops. Thus, purified TloR directly cleaves tRNA and preferentially targets the anticodon loop. The ability to cleave an unmodified *in vitro*-transcribed substrate further demonstrates that native tRNA modifications are not required for activity.

The predominant fragment endpoints obtained from *E. coli* did not precisely coincide with the direct cleavage pattern observed *in vitro*. Whereas purified TloR cleaved predominantly within the anticodon loop, TloR-enriched 5′ fragments isolated from cells frequently terminated further upstream and formed ladders of closely spaced endpoints (Fig. 4A). Because small-RNA sequencing captures fragments that accumulate after intoxication rather than primary cleavage events alone, these differences are consistent with secondary processing of TloR-generated products by endogenous RNases (Fig. 4F) [26]. Together, the cellular and biochemical data establish TloR as a broadly acting tRNase that targets diverse tRNA substrates while preferentially cleaving the anticodon loop.

## DISCUSSION

Most characterized bacterial tRNase toxins achieve specificity by discriminating among individual tRNAs or restricted substrate subsets [20, 27–29]. TloR reveals a different targeting strategy. Rather than exhibiting strong tRNA-isotype specificity, TloR acts across tRNAs decoding 19 of the 20 amino acids while retaining a pronounced positional preference for exposed regions of the tRNA, particularly the anticodon loop. TloR therefore appears to shift specificity from substrate identity toward substrate architecture: recognition is broad across the tRNA pool, but cleavage remains positionally constrained. Cleavage within the tRNA pool is sufficient to arrest translation, because purified TloR blocked reporter protein synthesis in a reconstituted system and toxin expression in cells produced no detectable loss of 16S or 23S rRNA. This combination distinguishes TloR from characterized tRNases such as colicin D and colicin E5 [4, 7, 8], which recognize restricted sets of tRNAs, and provides a different solution for disrupting translation through tRNA cleavage.

Neither broad RNA cleavage nor anticodon-loop targeting is unprecedented on its own, but we are not aware of a characterized toxin that combines them. Cleavage within the anticodon loop is a recurring strategy among RNA-targeting toxins, used by colicin E5, the phage-activated anticodon nuclease PrrC, and several VapC toxins, in each case against a single tRNA or a small set of isoacceptors [7, 28, 29]. In contrast, general tRNA degradation has so far been only reported for enzymes whose substrate range extends well beyond tRNA. The recently characterized polymorphic toxin YxiD was described as a tRNase because it cleaved tRNAs, but it also degraded additional RNA substrates [30]. Our structure-guided classification places YxiD near the

RNaseA/Ntox50 branch of canonical BECR proteins, a group associated with broad unspecific RNase activity [14]. YxiD therefore supports broad RNA cleavage rather than selective attack on the tRNA pool. Similarly, CdiA-CT^O32:H37^ and TseR degrade a wide range of tRNA, but respectively also degrade 16s rRNA or single-stranded RNA indiscriminately [31, 32]. TloR differs in that the cleavage products identified here derive from tRNAs, and direct cleavage of a defined tRNA occurs preferentially within the anticodon loop. The distinguishing feature of TloR is the combination of broad tRNA targeting with positional bias.

The molecular basis for this broad recognition remains to be determined, but several observations suggest that TloR recognizes properties shared among folded tRNAs. TloR-associated fragments arise from sequence-divergent tRNAs but exhibit similar positional distributions, and purified TloR cleaves an *in vitro*-transcribed tRNA lacking native modifications. These observations argue against an absolute requirement for a particular modification in the substrate tested and are consistent with recognition of structural or physicochemical features common to tRNAs. Such features could include the geometry of exposed loops, accessibility of the phosphodiester backbone, or electrostatic properties of the folded RNA. The preference for the anticodon loop may therefore reflect its accessibility within the tRNA architecture.

TloR also expands the emerging functional diversity of the recently described BECR-like superfamily [14]. The other experimentally characterized BECR-like RNase, SRLase1, recognizes a fundamentally different substrate context: it targets the sarcin-ricin loop of 23S rRNA through an EF-Tu-dependent mechanism [15]. TloR instead directly attacks tRNA and does not require native tRNA modifications for cleavage. These contrasting activities suggest that the BECR-like catalytic architecture can support substantial diversification in substrate recognition while retaining an RNase function. The requirement for His68 and His79 supports the structure-based inference that BECR-like proteins retain the catalytic organization of canonical BECR ribonucleases, which cleave RNA by metal-independent transesterification and leave 2′,3′-cyclic phosphate and 5′-hydroxyl termini [7, 13]. This interpretation parallels the broader evolutionary plasticity associated with canonical BECR proteins and suggests that the divergent BECR-like superfamily may contain additional RNA-targeting activities that cannot be inferred readily from sequence.

The predicted TloR-TloRI complex further suggests a simple mechanism for immunity that takes advantage of the same surface used for substrate recognition. TloR contains a positively charged pocket, while TloRI presents a complementary negatively charged surface across this region. Mutations in TloR residues adjacent to the putative active site can disrupt neutralization without eliminating toxin activity. We therefore propose that TloRI acts as a substrate mimic, using its negatively charged surface to occupy the RNA-binding region and prevent substrate access to the active site. Electrostatic occlusion is one of two main recurring solutions to the immunity problem. Barstar, the inhibitor of the canonical BECR barnase, blocks the barnase active site directly, with two of its acidic residues occupying the phosphate-binding subsites used by RNA [33, 34]. The immunity proteins of the nuclease colicins instead bind an exosite adjacent to the catalytic center and inhibit indirectly [35]. The predicted TloR-TloRI arrangement falls in the first class, as do several other characterized toxin-immunity pairs [13, 24, 25, 36, 37].

The broad substrate range of TloR may also have consequences for interbacterial competition. TloR homologs occur across diverse bacterial lineages and are repeatedly associated with several antagonistic systems. By targeting many tRNAs rather than relying on recognition of a small substrate subset, TloR activity may be comparatively robust to variation in individual tRNA sequences. Recognition of features shared among folded tRNAs would also be expected to extend to phylogenetically distant prey, which is consistent with the recurrent recruitment of TloR by several secretion systems in bacteria that occupy very different niches. Because the producing cell is protected by a dedicated immunity protein rather than by altering its own tRNAs, breadth of substrate range imposes little cost on the attacker. This possibility is particularly interesting because variation in anticodon-loop sequences of phage-encoded tRNAs can confer resistance to host anticodon nucleases [38], demonstrating that tRNA diversification can alter susceptibility to RNA-targeting defenses. Additionally, bacterial RNA repair systems, including RtcB and the Pnkp-Hen1 cassette present in other lineages, reseal tRNAs broken within the anticodon loop by ribotoxins [39, 40]. Whether prey RNA repair limits TloR-mediated killing remains to be tested, but broader tRNA cleavage creates a larger burden on prey cells to rapidly repair RNA.

Together, this work identifies a mode of tRNase activity in which specificity is given by structural context. TloR combines broad recognition of the tRNA pool with preferential cleavage of exposed tRNA loops, extending the functional diversity of BECR-like ribonucleases and revealing how an antibacterial toxin can achieve substrate breadth without abandoning positional specificity. Identifying the structural determinants that TloR recognizes will show how general this strategy proves to be among antibacterial ribonucleases.

## METHODS

### Strains and plasmids

Bacterial strains, DNA oligonucleotides, plasmids, and primers are listed in Table S3. Strains were grown at 37 °C with agitation in lysogeny broth containing 10 g/L tryptone, 10 g/L NaCl, and 5 g/L yeast extract. AB medium used for the SOS-response assay contained 0.2% (NH4)_2_SO_4_, 0.6% Na_2_HPO_4_, 0.3% KH_2_PO_4_, 0.3% NaCl, 0.1 mM CaCl_2_, 1 mM MgCl_2_, and 3 µM FeCl_3_, supplemented with 0.2% sucrose, 0.2% casamino acids, 10 µg/mL thiamine, and 25 µg/mL uracil.

TloR was cloned into pBRA vector under the control of the P_BAD_ promoter [41]. TloRI was cloned into pEXT22 vector under control of the P_TAC_ promoter [42]. For protein expression and purification, TloR was cloned into pRSFDuet including an N-terminal 6xHis tag and C-terminal Strep II tag. Point mutations were created via splicing by overlap extension (SOE) PCR [43]. All constructs were confirmed by sequencing. Chromosomal deletions in *S.* Oranienburg were constructed using λ-Red recombineering [44]. For complementation of Δ*tssL* the pFPV25.1 plasmid was used [45].

### Bioinformatic analysis

Bioinformatic analysis was performed as previously described [36]. Sequences of TloR and TloRI from *S. enterica* serovar Oranienburg were used as queries for iterative jackhmmer [46] searches against the RefSeq [47] and UniProt [48] databases. Searches were performed until exhaustion, meaning until retrieval of additional homologs stopped. Homologs were manually curated to retain preservation of secondary structure and aligned using MAFFT [49] and refined in Jalview Version 2.11.5.1 [50]. Amino acid sequence logos were generated using WebLogo [51, 52]. Taxonomic distribution was visualized using AnnoTree [53]. Genomic contexts were visualized as described previously [36]. In short, sequence alignments were converted to profile hidden Markov models and queried against the NCBI protein database [54] using HMMER [55]. Homologs were retrieved using an E-value cutoff of 1 x 10^-25^. Gene neighborhoods were collected and visualized using the ROTIFER package (https://github.com/leepusp/rotifer) with default parameters. Sequence similarity network was generated on the EFI-EST platform [56] using the organic layout from yFiles [57] and visualized in Cytoscape [58].

### Toxicity assay

*E. coli* cells were co-transformed with plasmid encoding for toxin (pBRA TloR) and immunity protein (pEXT22 TloRI) and selected on LB agar with 50 μg/mL streptomycin, 50 μg/mL kanamycin, and 0.2% D-glucose. Colonies were inoculated in LB media overnight at 37 °C with the same antibiotics. Cells were adjusted to OD_600_ = 1.0, serially diluted four-fold, and 4.5 µL was spotted onto LB agar containing either 0.5% D-glucose or 0.2% L-arabinose plus 200 μM IPTG, along with the same antibiotics. The plates were then incubated at 37 °C overnight and subsequently imaged. All assays were performed in three independent biological replicates.

### Interbacterial competition

Interbacterial competition assays were performed using *S. enterica* serovar Oranienburg wild-type or *ΔtssL* as attackers and *S.* Oranienburg *ΔtloR*/*tloRI*::Cm as prey. The Δ*tssL* mutant was complemented with *tssL* using pFPV25.1 [45]. Overnight cultures were diluted 1:100 in LB and grown to an OD_600_ of 0.6 to 0.8. Attacker and prey cultures were adjusted to OD_600_ values of 4.0 and 1.0, respectively, and mixed in a 1:1 volume ratio. Mixtures were spotted onto a 0.22 μm nitrocellulose membrane placed on LB agar plates and incubated for 4 h at 37 °C. Bacteria were resuspended in LB, serially diluted, and plated on media selective for the prey strain. Prey recovery rate was calculated as the ratio of output of colony-forming units (CFU) to input CFU and normalized to wild-type values.

### SOS response

*E. coli* expressing pSC101-P*recA*::GFP [59] and TloR (WT or H79A) were subcultured in streptomycin, kanamycin, and 0.2% glucose to OD_600_ = 0.6. Cultures were mixed with AB medium (1:1), AB medium with 0.2% D-glucose, or AB medium with 0.2% L-arabinose and homogenized in a 96-well plate. Bacterial growth (OD_600_) and GFP signal were measured every 15 min for 6 h at 37 °C using a BioTek Synergy H1 Multimode Reader. DNase TseV2 was used as a positive control [25] and PBS was used as a negative control.

### Plasmid DNA cleavage

*In vivo* plasmid degradation assay. *E. coli* carrying empty vector or TloR_WT_ expression constructs were grown to OD_600_ = 0.6 and induced or repressed for 1-2 h. Plasmid DNA was purified using the GeneJET Plasmid Miniprep Kit and analyzed by electrophoresis on 1% agarose gels containing SYBR Safe. DNA was visualized using a ChemiDoc imaging system. *In vitro* plasmid degradation assay. Plasmid DNA was incubated alone, with purified DNase Rehh [36], TloR_WT_, or TloR_H79A_ for 1 h. DNA was analyzed by electrophoresis on 1% agarose gels containing SYBR Safe. DNA was visualized using a ChemiDoc imaging system.

### Total RNA cleavage

*E. coli* cells expressing TloR (WT or H79A) were subcultured to OD_600_ = 1.0 in LB with streptomycin and 0.5% D-glucose. Cultures were spun down and resuspended in LB with streptomycin and 0.2% L-arabinose. Induced cultures were incubated at 37 °C for 15 min and RNA was extracted using TRIzol (Invitrogen). RNA was denatured for 2 min at 70 °C and separated on a 1.5% agarose gel. RNA was stained with SYBR Gold (1:10,000) for 30 min and visualized on a Bio-Rad ChemiDoc imaging system. To resolve small RNAs from the same extracts, RNA was separated in parallel on an 8 M urea, 10% polyacrylamide gel, stained with SYBR Gold, and imaged on the same system.

### Protein expression and purification

*E. coli* BL21(DE3) carrying pRSFDuet 6xHis-TloR_WT_-Strep or 6xHis-TloR_H79A_-Strep were grown in 1L of LB supplemented with kanamycin and 0.5% glucose until OD_600_ = 0.6. Expression was induced with 1 mM IPTG for 4 h at 37 °C. Cells were harvested via centrifugation at 10,000 *g* for 15 min at 4 °C, and pellets were resuspended in lysis buffer (100 mM Tris-HCl pH 7.5, 500 mM NaCl, 5 mg/mL lysozyme, 5% glycerol) and lysed at 4°C through 7 cycles (15 sec ON/25 sec OFF) at 50% amplitude with a Qsonica Q700 sonicator. The lysate was centrifuged at 35,000 *g* for 40 min at 4°C. Supernatant was filtered using 0.22 μm PES filter and loaded onto a 1 mL StrepTrap HP column (Cytiva) equilibrated in wash buffer (100 mM Tris-HCl pH 7.5, 500 mM NaCl, 5% glycerol). The column was washed with 30 column volumes (CV) of wash buffer, then eluted with 7 CV of elution buffer (100 mM Tris-HCl pH 7.5; 500 mM NaCl; 5% glycerol; 50 mM biotin). 1 mL fractions were separated on 15% SDS-PAGE and probed via western blot to detect fractions containing TloR protein. Fractions were combined and loaded onto a 1mL HisTrap HP (Cytiva) equilibrated in wash buffer (50 mM Tris-HCl pH 7.5, 350 mM NaCl, 10 mM imidazole, 5% glycerol, 5 mM β-mercaptoethanol). The column was then washed with 10 CV of wash buffer, followed by a 20 CV gradient of guanidinium hydrochloride in wash buffer up to a maximum of 6 M. Five additional CV of 6 M guanidinium hydrochloride was used to fully denature TloR and remove the immunity protein TloRI. The column was then washed with a 20 CV downward gradient for refolding, followed by 10 CV of wash buffer. The column was eluted in 20 CV of a gradient of elution buffer (50 mM Tris-HCl pH 7.5, 500 mM imidazole, 350 mM NaCl). Fractions were buffer exchanged (100 mM Tris-HCl pH 7.5, 150 mM NaCl, 5% glycerol) and concentrated using an Amicon of 30 kDa (Sigma). Protein aliquots were snap-frozen until use.

### *In vitro* translation Assay

*In vitro* protein synthesis was performed with the PURExpress In Vitro Protein Synthesis Kit (NEB). Each reaction contained 10 µL solution A, 7.5 µL solution B, 250 ng DHFR-FLAG template DNA, and 20 units of murine RNase inhibitor (NEB, M0314), according to the manufacturer’s instructions. Reactions received 200 nM of the recombinant protein, or the corresponding buffer control, and were incubated for 4 h at 37 °C. Reactions were stopped on ice. Protein from 4 µL of reaction was separated on a 15% SDS-PAGE, transferred to a nitrocellulose membrane and incubated with polyclonal anti-FLAG antibody, followed by secondary antibody. Signals were detected with ECL substrate and visualized on a Bio-Rad ChemiDoc imaging system. Anti-EF-Tu was used as a loading control.

### RNA-seq

Small-RNA sequencing was performed by CD Genomics using its tRF and tiRNA sequencing workflow. *E. coli* MG1655 cells carrying TloR or empty-vector constructs were grown to OD_600_ = 1.0, and total RNA was extracted with TRIzol. RNA terminal modifications were treated (3’ deacylation, 3’-cP removal, 5’ phosphorylation, and m1A/m3C demethylation) to improve adapter ligation. Adapters were ligated to the 3’ and 5’ ends, cDNA was synthesized and PCR-amplified, and library products corresponding to 14-to 70 nt RNA inserts were size-selected. Libraries were sequenced on an Illumina platform. Three biological replicates were analyzed for each condition. Reads were mapped to mature and precursor tRNA allowing 1 bp mismatch, and sequencing counts were normalized by counts per million (CPM) of total aligned reads. Fold change and log2 fold change (log2FC) were calculated by comparing TloR and control CPM. To create the heat map and tRNA histograms, reads were filtered by mature tRNA and selected only for reads in which the start or end position was < 4 nucleotides from the corresponding end of the tRNA using R. Log2FC was plotted on the heat map by tRNA species and 3’ end position. Histograms were plotted as log2FC over the nucleotide position of the other end of the tRNA fragment. For aligned reads, log2FC was multiplied by 10 to capture decimal places, then a random, representative subset of 25% of the data was taken. This was plotted according to nucleotide position, with lines being proportionally representative of the log2 fold increase of their corresponding fragments.

### *In vitro* tRNA synthesis and cleavage

tRNA was synthesized via *in vitro* transcription (IVT). Single-stranded DNA oligos encoding tRNA with an upstream T7 promoter sequence were ordered from IDT and used as templates for PCR. To synthesize tRNA, 0.5 μg of PCR product was used as a template for an IVT reaction using the HiScribe T7 High Yield RNA Synthesis kit (NEB, E2040). The 30 μL reactions were prepared following the manufacturer’s instructions and incubated for 16–18 h at 37 °C. IVT products were treated with DNase I (NEB, M0303) and then purified using RNA Cleanup kit (NEB, T2050). For *in vitro* cleavage, 100 nM of recombinant protein were incubated with 100 nM of tRNA in reaction buffer (20 mM Tris-HCl pH 7.5, 10 mM MgCl_2_, 100 mM NaCl). All reactions were incubated for 1 h at 37 °C and mixed 1:1 with 2x RNA loading dye to stop the reaction. Samples were incubated for 2 min at 70 °C before loading onto a 15% urea–PAGE. The gel was run at 25 mA, stained with SYBR Gold for 30 min and imaged on a Bio-Rad ChemiDoc.

### Statistical analysis

All statistical analysis was performed in GraphPad Prism. Statistical tests used are indicated in corresponding figure legends. For interbacterial competition assays, data represents the mean ± SD from five independent experiments.

## ACKNOWLEDGMENTS

This research was supported by startup funds from UT Austin College of Natural Sciences at The University of Texas at Austin to E.B.-S. The authors acknowledge the Texas Advanced Computing Center (TACC) at The University of Texas at Austin for providing computational resources that have contributed to the research reported in this study (http://www.tacc.utexas.edu). Figures were created in BioRender. Lin, J. (2026) https://BioRender.com/bcxechs. Microsoft Copilot was used to assist with language editing of the manuscript. The authors reviewed and edited all text and take full responsibility for the content of this publication.

## SUPPLEMENTARY LEGENDS

**Fig S1. TloR is associated with several diverse immunity families. (a)** Sequence similarity network of TloR Immunities. EFI-EST was used to produce a sequence similarity network of the immunity homologs of TloR. Edges were drawn with e-value threshold of 1e-1 and the organic layout from the yFiles was used. TloR Immunities annotated as Imm55 cluster independently with previously described Imm55 proteins with some intersection. Imm70 immunities cluster independently from either family. **(b)** TloR homologs show limited pairwise sequence alignment but retain a highly similar structural fold. In comparison, associated immunity proteins show both limited pairwise sequence alignment and limited secondary structure similarity. **(c)** Associated immunities Imm55.2 and Imm70 bind to TloR with a high predicted interaction score. Structure-based alignments were performed using Mol* [60] and figures were generated with ChimeraX [61].

**Fig S2**. TloR point mutants expression control. **(a)** Western blot of TloR mutants. Point mutants are stable enough for protein to be sufficiently accumulated and probed. TloR_WT_ is not detected due to toxicity. DnaK was used as a loading control. **(b)** TloR mutants toxicity assay. Serial dilutions of *E. coli* carrying wild-type or point mutants of TloR plated in repressive or inductive conditions. Images are representative of three independent experiments.

**Fig S3. Recombinant protein expression and purification.** TloR_WT_ **(a)** and TloR_H79A_ **(b)** protein purification. SDS-PAGE following initial Strep-tagged purification (left) and subsequent 6xHis-tag purification (right). Soluble (SOL) and insoluble (INS) fraction of lysate, flow through (FT), wash, and elutions. **(c)** SDS-PAGE of purified proteins used for assays.

**Fig S4.** RNA-seq data shows widespread tRNA cleavage. Sequencing reads mapping to representative substrates. Read abundance is represented as the log2FC in TloR-expressing cells relative to the control. Dotted lines delimit the anticodon loop.

**Table S1. List of species encoding TloR homologs.**

**Table S2. TloR tRF and tiRNA RNA-sequencing raw data.**

**Table S3. List of primers, DNA oligos, plasmids, and strains used in the study.**

**Dataset S1. TloR homologs and their genomic context.**

