## Supplementary material for "An RNase bacterial toxin family cleaves the anticodon loop across diverse tRNA isotypes": Fig. S

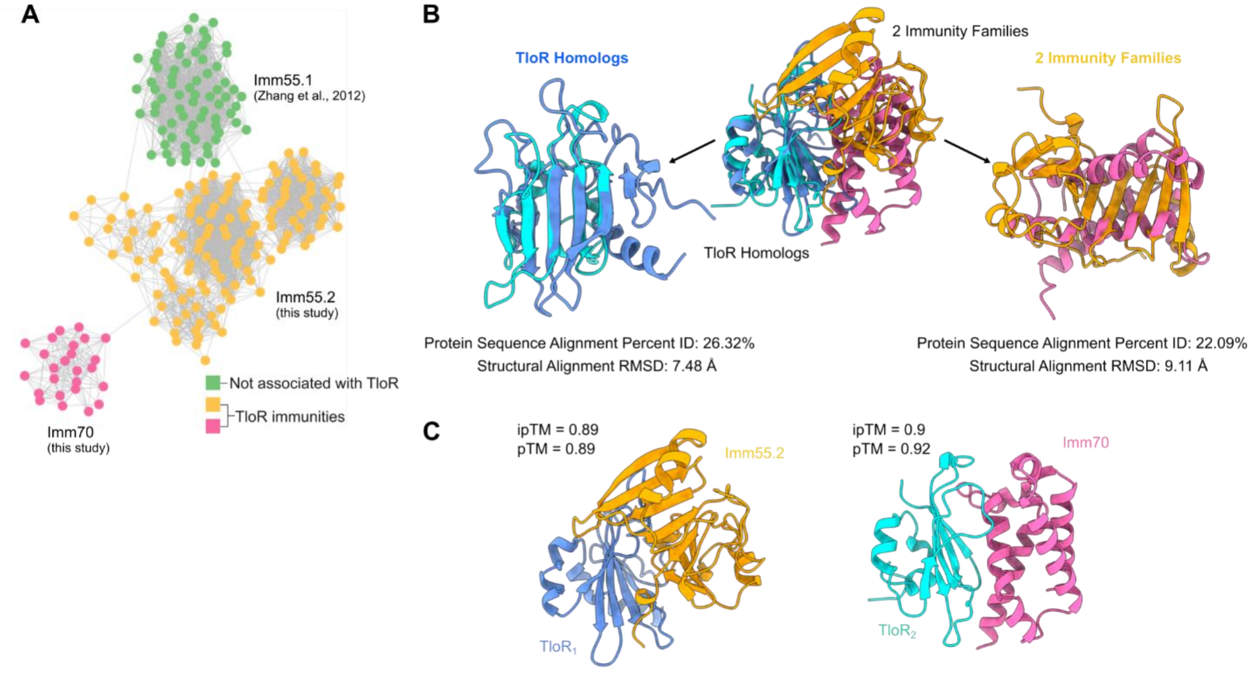


**Fig S1. TloR is associated with several diverse immunity families. (a)** Sequence similarity network of TloR immunity proteins. EFI-EST was used to produce a sequence similarity network of the immunity protein homologs of TloR. Edges were drawn with e-value threshold of 1e-1 and the organic layout from the yFiles was used. TloR immunity proteins annotated as Imm55 cluster independently with previously described Imm55 proteins with some intersection. Imm70 immunities cluster independently from either family. **(b)** TloR homologs show limited pairwise sequence alignment but retain a highly similar structural fold. In comparison, associated immunity proteins show both limited pairwise sequence alignment and limited secondary structure similarity. **(c)** Associated immunities Imm55.2 and Imm70 bind to TloR with a high predicted interaction score. Structure-based alignments were performed using Mol* [60] and figures were generated with ChimeraX [61].

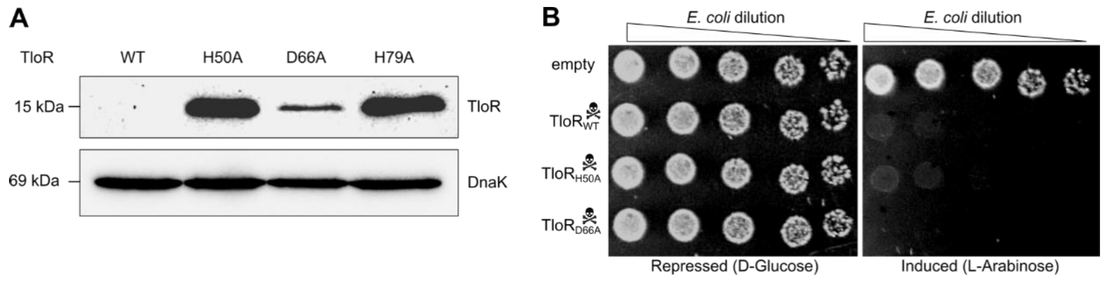


**Fig S2. TloR point mutants expression control. (a)** Western blot of TloR mutants. Point mutants are stable enough for protein to be sufficiently accumulated and probed. $TloR_{WT}$ is not detected due to toxicity. DnaK was used as a loading control. **(b)** TloR mutants toxicity assay. Serial dilutions of *E. coli* carrying wild-type or point mutants of TloR plated in repressive or inductive conditions. Images are representative of three independent experiments.

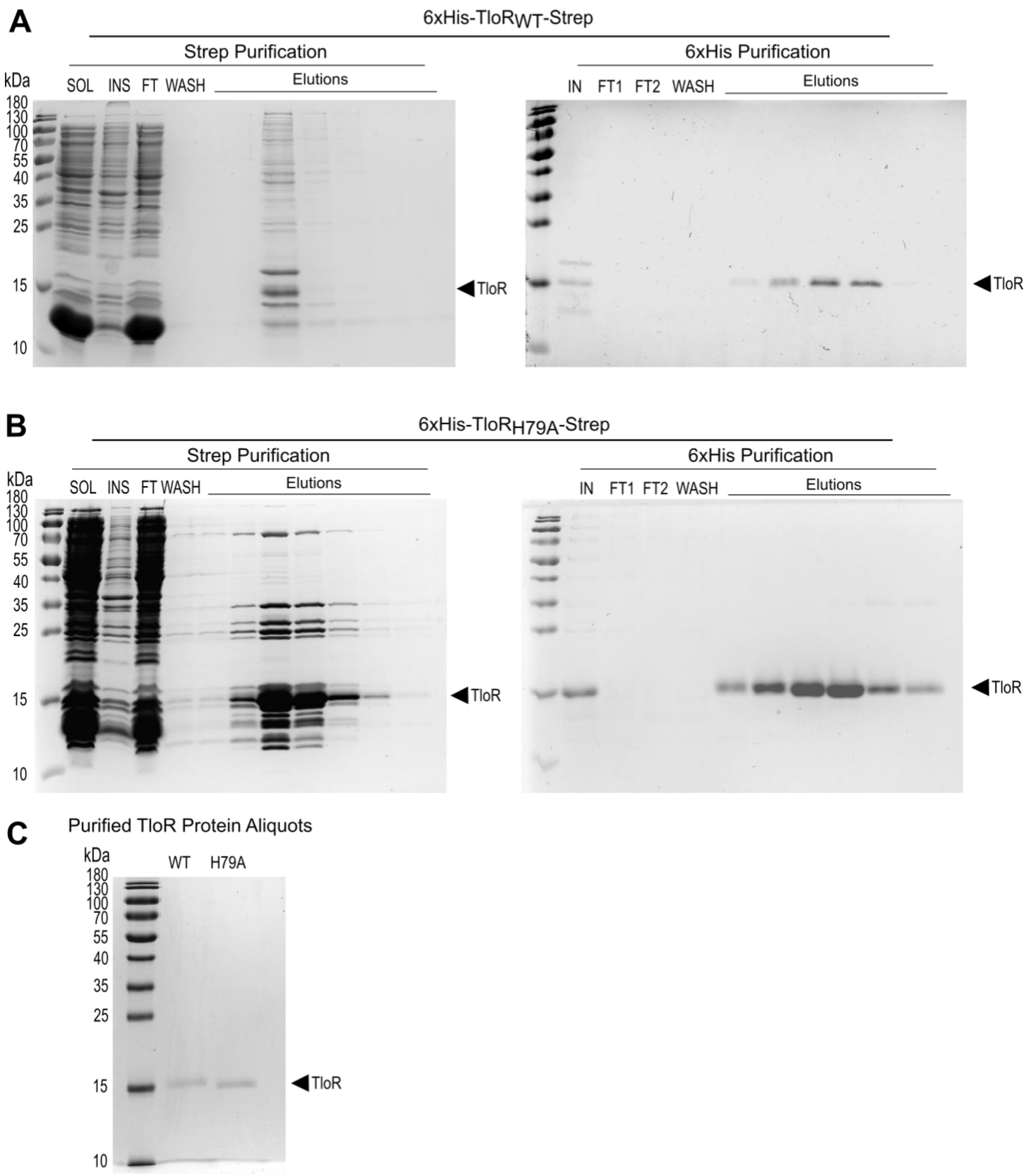


**Fig S3. Recombinant protein expression and purification.** TloR$_{WT}$ **(a)** and TloR$_{H79A}$ **(b)** protein purification. SDS-PAGE following initial Strep-tagged purification (left) and subsequent 6xHis-tag purification (right). Soluble (SOL) and insoluble (INS) fraction of lysate, flow through (FT), wash, and elutions. **(c)** SDS-PAGE of purified proteins used for assays.

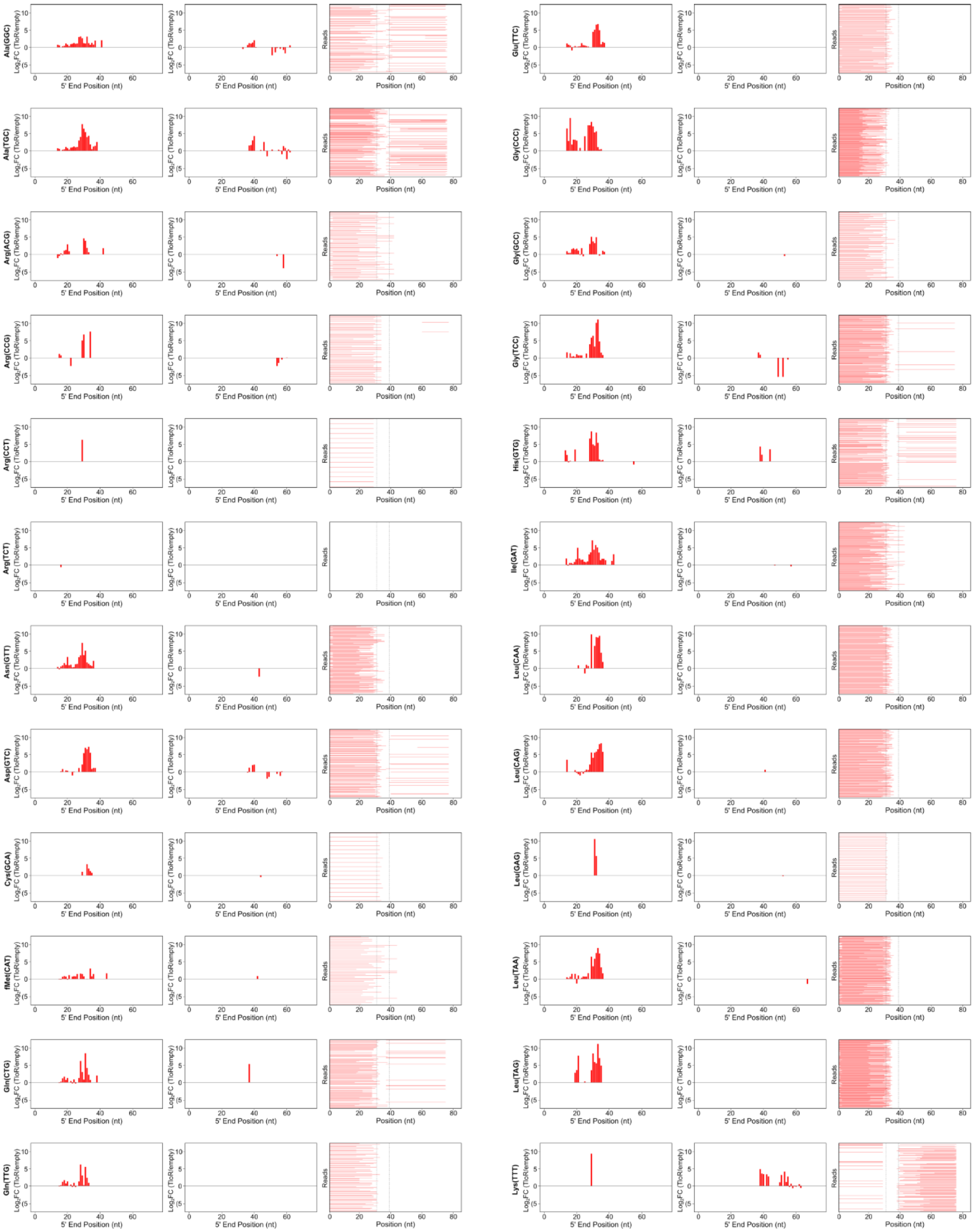
Ala(GGC)
Ala(TGC)
Arg(ACG)
Arg(CCG)
Arg(CCT)
Arg(TCT)
Asn(GTT)
Asp(GTC)
Cys(GCA)
fMet(CAT)
Gln(CTG)
Gln(TTG)
Glu(TTC)
Gly(CCC)
Gly(GCC)
Gly(TCC)
His(GTG)
Ile(GAT)
Leu(CAA)
Leu(CAG)
Leu(GAG)
Leu(TAA)
Leu(TAG)
Lys(TTT)
Log$_2$FC (TloR/empty)
5' End Position (nt)
Reads
Position (nt)

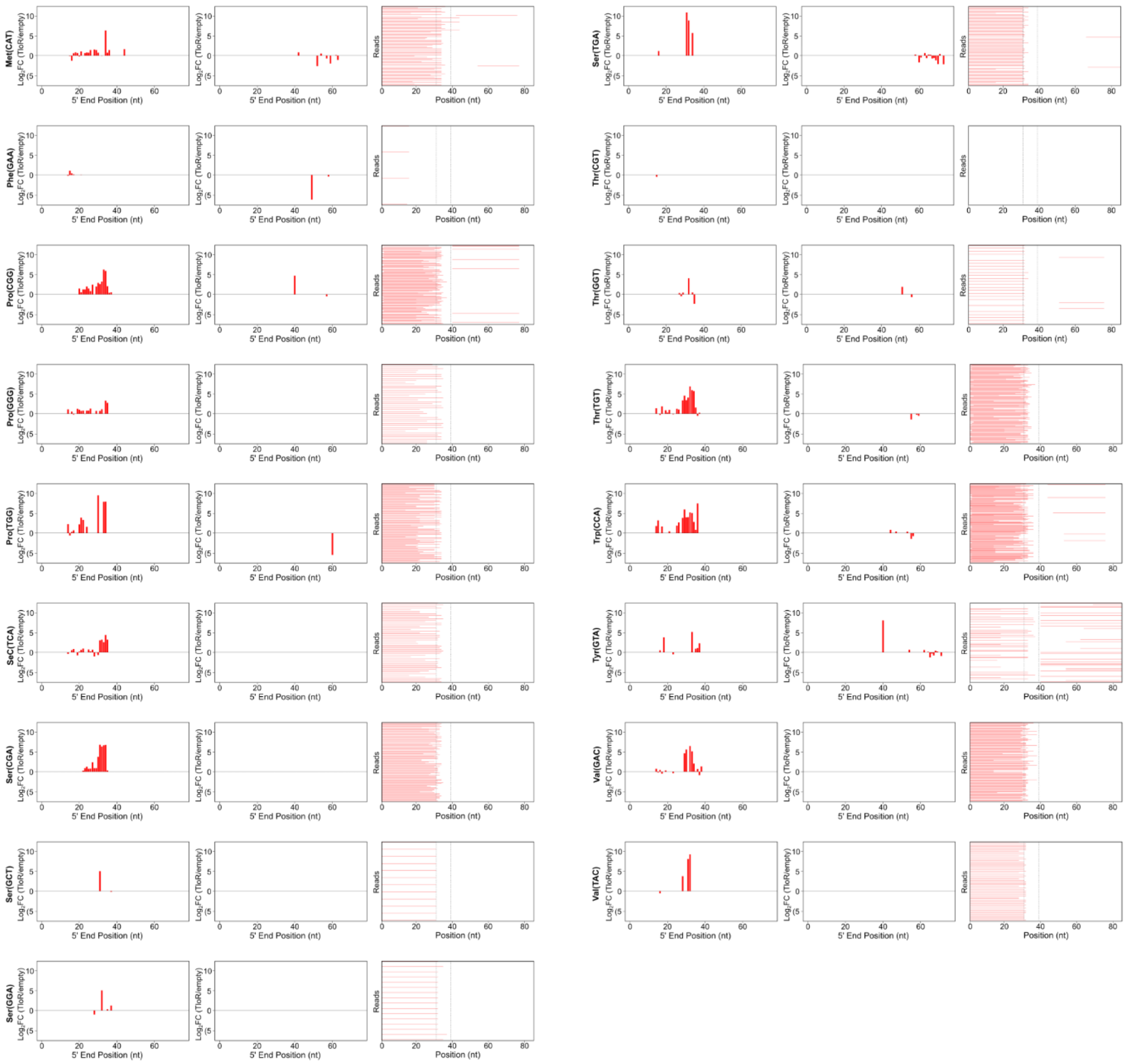


**Fig S4.** RNA-seq data shows widespread tRNA cleavage. Sequencing reads mapping to representative substrates. Read abundance is represented as the log2FC in TloR-expressing cells relative to the control. Dotted lines delimit the anticodon loop.
